# Planning and scheduling biological experiments across multiple liquid handling robots

**DOI:** 10.64898/2025.12.26.696584

**Authors:** Benjamin M. David, Paul A. Jensen

**Author notes:** Correspondence to. Website: http://jensenlab.net.

## Abstract

Coordinating multiple liquid handling robots is a complex logistical task when designing biological experiments. Protocol designers must consider the capabilities and constraints of each robot to distribute work optimally across multiple instruments. We developed an optimization framework that finds optimal liquid handling solutions that leverage an arbitrary number of robots. Our algorithm, called Pourfecto, abstracts the capabilities of each robot and their labware, allowing us to plan and schedule a wide range of biological experiments using commercial instruments and custom-built hardware. Pourfecto can optimize multiple objectives (minimum transfers, fewest reagents, fewest labware swaps) and scales to experiments with hundreds of thousands of liquid transfers.

## 1 Introduction

Liquid handling robots are the backbone of high-throughput laboratory sciences. Platforms that combine multiple liquid handlers power drug screening campaigns [1], clinical diagnostics [2], chemical synthesis pipelines [3], [4], and robot scientists that increase the pace of science [5]–[8].

Scientists and engineers who adopt automation must translate their experiments into protocols for the instruments. Protocol design can be broken down into planning and scheduling phases [9]. A liquid handling plan determines the necessary steps to transform the available reagents into the target solutions for an experiment. For example, a plan might include mixing 10 *μ*l of stock A and 30 *μ*l from stock B to create target stock C. Planning ensures that all transfers in the protocol use available reagents, that sufficient quantities of the reagents are available, and that the final solutions match the targets specified by the user.

The scheduling phase determines how liquid handlers will execute the plan[10]. Scheduling considers all the capabilities and constraints of the robots, including the space available on the decks, the volumes of labware, and the minimum and maximum volumes that can be dispensed by each instrument. Liquid handlers are often specialized for particular tasks, such as bulk filling labware, mixing combinations of reagents, or stamping liquid from one location to another. An optimal schedule will assign operations to each instrument to maximize efficiency, improve accuracy, or decrease runtimes and costs.

The liquid handling industry continues to grow and specialize, and tools for planning and scheduling protocols must become increasingly flexible to capitalize on the latest hardware developments. Most liquid handler software is designed to be used by humans. These tools generate machine-readable instructions from human-designed protocols and provide graphical interfaces to map out laboratory workflows [11]– [14]. In the era of AI-driven science, planning and scheduling are performed by model-driven software that benefits from a programmatic interface. Autonomous systems cannot rely on humans to select the best set of instruments to complete a task. Prior work on liquid handler planning and scheduling considers protocols executed on a single robot [15], but true autonomy requires software that optimally schedules across multiple instruments.

This work solves the problem of planning and scheduling experiments across a diverse set of liquid handling robots. We show how planning and scheduling can be unified under a single combinatorial optimization framework, and we introduce the Pourfecto algorithm to plan and schedule multiple liquid handlers on demand. Pourfecto first plans by searching for a feasible set of liquid transfers that combine source stocks into the desired target stocks. Second, Pourfecto schedules by assigning transfer operations to robots based on their capabilities, constraints, and user-defined costs. The result is a single optimal protocol that satisfies users’ experimental (planning) and operational (scheduling) goals. The Pourfecto framework allows users to define and minimize operational costs, and the framework can be extended to any liquid handling instrument, from manual pipettes to high-throughput robots. We demonstrate how Pourfecto can plan and schedule protocols that execute thousands of liquid transfers across multiple instruments in a single experiment.

## 2 Methods

### 2.1 Planning view formalism

Liquid handling is simultaneously a planning and scheduling problem. The planning viewpoint considers a set of source stocks that must be combined by liquid transfers to create a set of target stocks (Figure 1a). Each stock is located in a “well” (e.g. a tube, reservoir, or well of a multiwell plate) and contains one or more reagents at a specified concentration (percent weight, mM, g/L, etc.). The planning objective is to find a set of well-to-well transfers that create the target stocks using the source stocks. Each transfer is a triplet of a source stock, a target stock, and the volume transferred from source to target. The planning problem is constrained by the concentrations and volumes of each source and target stock. Planning requires that no reagent is “overdrafted” by, for example, ensuring that 1.1 mL of reagent is not drawn from a source stock with a volume of only 1.0 mL. The target stocks must be made with the correct volume and concentration, which forces Pourfecto to use only the source stocks that contain a subset of the reagents in the target stock (Supplementary Material 1). (A source stock containing both glucose and fructose cannot be used to make a target stock that contains glucose but no fructose.)

**Figure 1:**
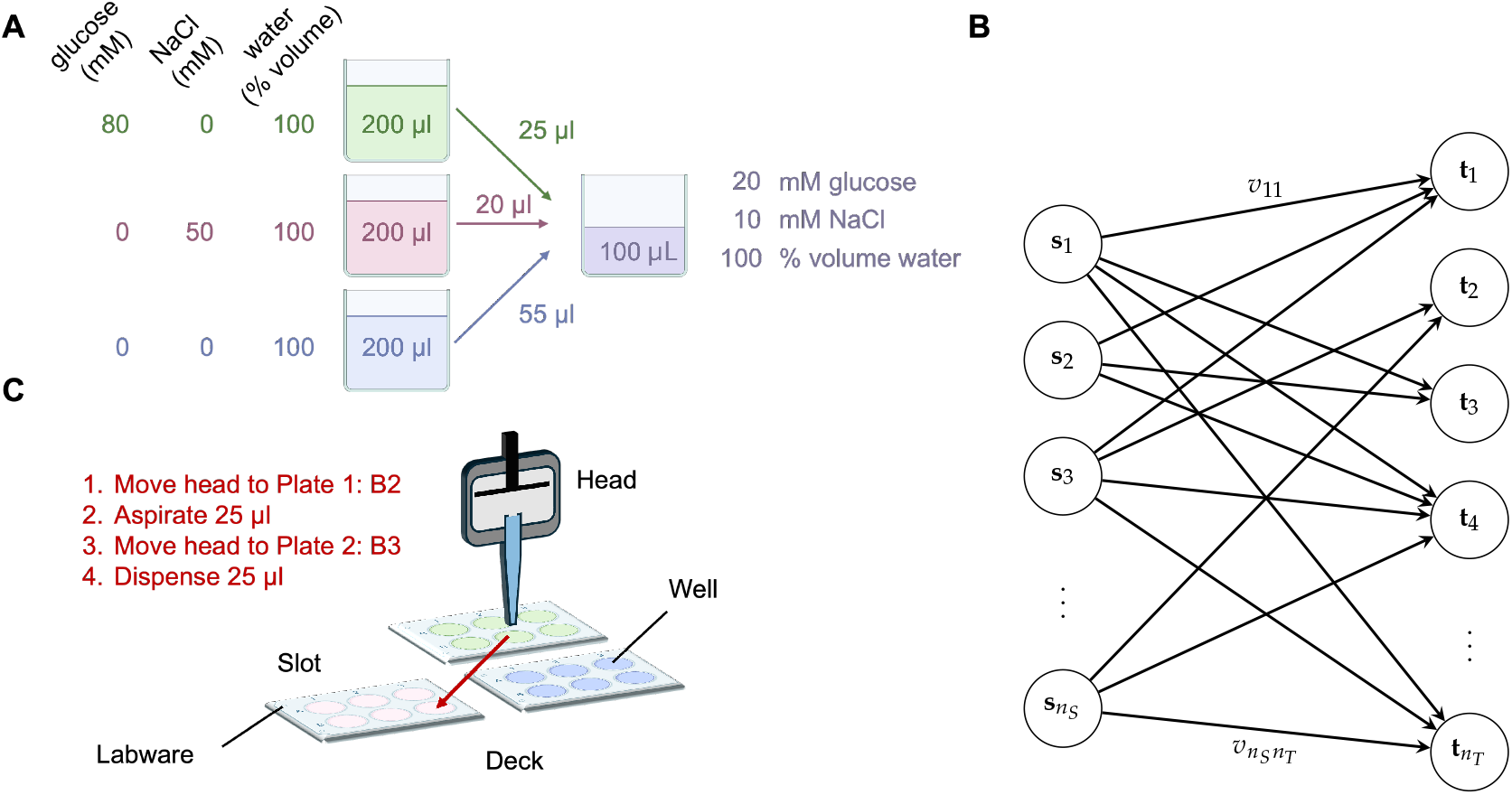
Optimal liquid handling protocols require planning and scheduling. **A**. Planning combines wells of source stocks to create the desired target stocks. Planning decisions generate well-to-well transfer volumes. **B**. Planning sets transfer variables (*v*_*st*_) on a bipartite graph connecting source stock wells **S** to target stock wells **T. C**. Scheduling determines how liquid handlers carry out the planned transfers. Optimal schedules execute the plan while satisfying the constraints of the liquid handlers and minimizing the cost of the operations.

Planning differs from the scheduling view in that planning disregards most of the logistics of moving reagents during the requested transfers. It is the scheduler, not the planner, that ensures the stocks are on the correct instrument, that the pipette head can reach the stocks, and that the pipette can hold the requested volume. The only decision variables during planning are the transfer volumes *v*_*st*_ that describe the volume of source stock *s* that should be transferred into target stock *t* (Figure 1b).

Planning calculates how transfers from *n*_*s*_ source stocks create *n*_*T*_ target stocks. Each stock includes a subset of the *n*_*R*_ reagents and is represented as a vector describing each reagent’s concentration. For example, consider a planning project with three reagents: glucose, sodium chloride (NaCl), and water. (Note that we include both solvents and solutes as reagents for liquid stocks.) Our reagents were used to make three 200 *μ*l source stocks

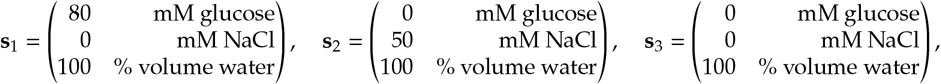

and our target is a 100 *μ*l stock of

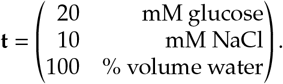

Pourfecto internally converts the units of the source and target stocks. The concentrations of reagents in stocks are converted to a quantity of reagent per unit of transferred stock. By default, Pourfecto calculates transfer volumes in *μ*l, so the units of glucose in stock **s**_1_ would be converted to 80 nmol/*μ*l. Target stocks are represented as quantities of reagent in the entire solution, so the units of glucose in **t** are 20 mM × 100 *μ*l = 2 *μ*mol. All of these unit conversions are handled automatically by Pourfecto, allowing users to specify the concentrations and volumes of the source and target stocks in convenient units.

A mass balance on each reagent constrains the possible transfer quantities to be

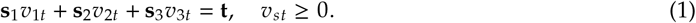

More generally, we can collect the *n*_*s*_ source stocks as columns of a source matrix **S** and rewrite Equation (1) as **Sv**_*t*_ = **t**, where 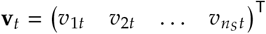 is a transfer volume vector that defines the volume transferred from each source stock *s* to make the target stock *t*. In the example above, the mass balance constraint requires that we transfer 25 *μ*l of stock 1, 20 *μ*l of stock 2, and 55 *μ*l of stock 3 to make the requested 100 *μ*l of the target stock (Figure 1a).

The mass balance constraint can be extended to multiple target stocks by collecting the target stock vectors into a matrix **T** and collecting the corresponding transfer volume vectors into a matrix **V**. The mass balance over all source and target stocks becomes the matrix linear system **SV** = **T**.

Mass balances are not the only planning constraints in practice. Each source stock has a finite volume, and users can limit the maximum volume of each target stock that can be made to avoid using too much reagent or overfilling the labware holding the target stock. The composition of the source stocks might also prevent us from making every target stock exactly. For example, source stocks of a low-solubility reagent might not be concentrated enough to make target solutions while leaving room to add other reagents. In such imperfect cases, our goal is to minimize the error between the stocks we make (**SV**) and the target stocks (**T**). Pourfecto, by default, minimizes the squared difference of this error and solves the constrained planning problem as the quadratic program

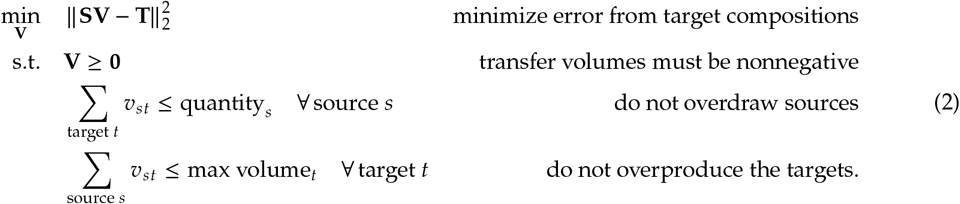

where ∥·∥ _2_ is the Frobenius norm, causing Pourfecto to search for plans that match the targets in the least-squares sense.

An optimal solution to the planning problem ensures that the transfers produce stocks that are as close as possible to the targets while satisfying practical constraints, such as source quantities and maximum target volumes. The logistical details of the transfers, however, require a detailed view of liquid handling that includes a mathematical definition of the instrument(s) completing the transfers. These logistical details are solved during the scheduling phase.

### 2.2 Scheduling view formalism

Liquid handlers execute three tasks: *aspirating* a liquid from a source well, *moving* that liquid to a new location, and *dispensing* it into the destination well (Figure 1c). Liquid handlers can be manually operated or robotic. A human-held pipette can adapt to many aspiration and dispensation volumes and move easily between locations. Robotic liquid handlers automate these tasks but vary tremendously in the labware they can access, the movements they can make, and the volumes they can aspirate and dispense.

We developed a general definition of liquid handlers that allows Pourfecto to schedule any manual or robotic liquid handler. To Pourfecto, a liquid handler combines two components: a *head* that holds pipette channels and pistons, and a *deck* that holds labware (wells) in slots.

The **head** contains channels that access wells and pistons that independently control aspiration and dispensation volumes^†^ (Figure 2a). A standard single-channel micropipette is the simplest robot, with a one-channel head connected to one piston. Handheld multichannel pipettes may have eight or twelve channels but only one piston, and aspirating 100 *μ*l with this piston will aspirate 100 *μ*l into each channel. Some robotic liquid handlers have multiple pistons connected to separate channels for independent control, e.g. an ARI Cobra (Figure 2a). The channels are typically arranged in a one-or two-dimensional array, with fixed spacing between channels. Multichannel instruments often space their channels to align with SBS/SLAS 96-well microplates [16]; however, some instruments use other spacings or have actuators that change the channel spacing on demand.

**Figure 2:**
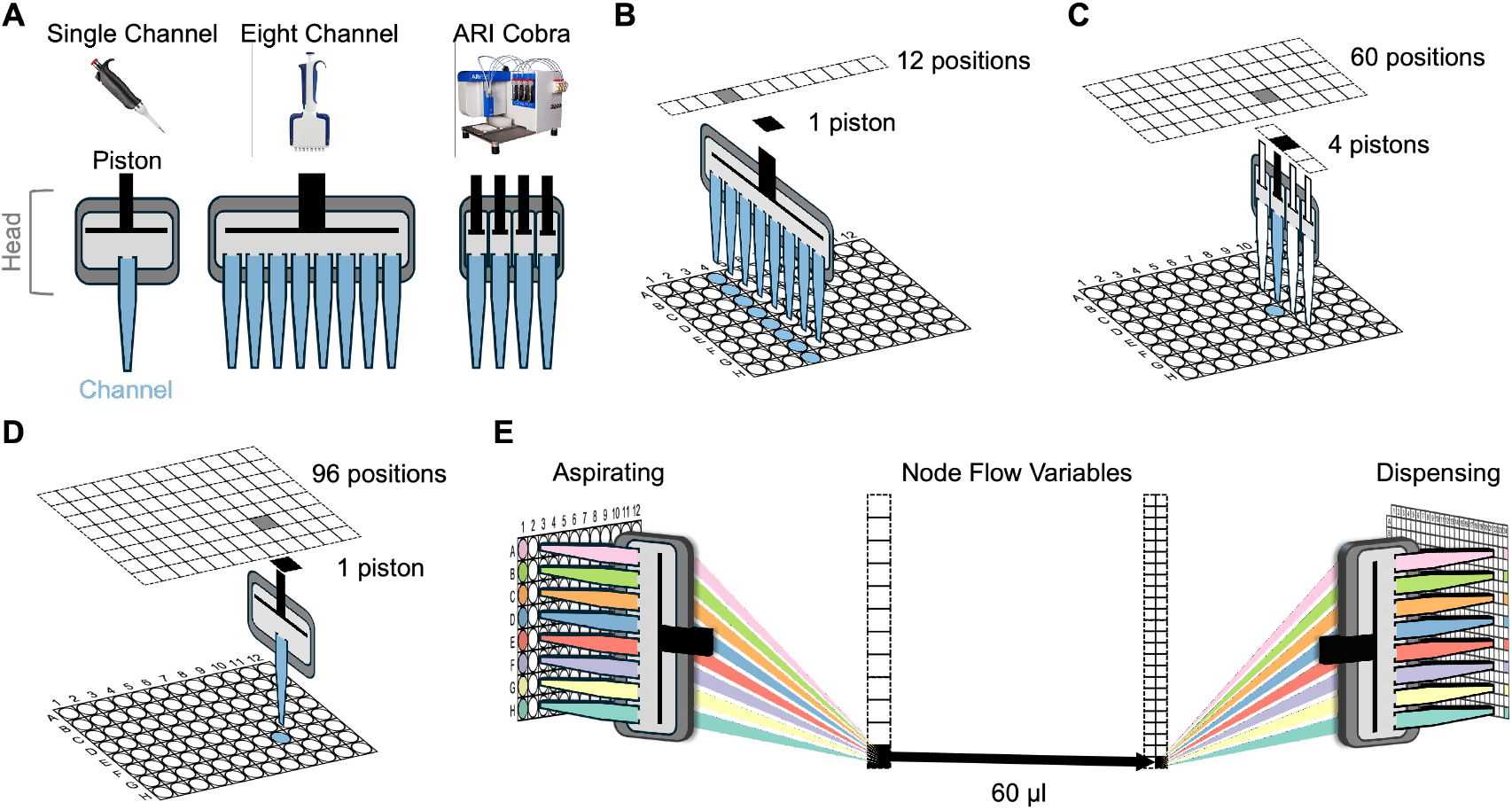
Liquid handlers include a pipetting head and a deck with slots for labware. **A**. The head is a set of interconnected pistons and channels. Pistons control aspiration and dispensation volumes, and channels connect the head to individual wells. **B-D**. Each head can access labware on the deck in a finite number of positions based on the head and labware geometries. Each piston/position pair connects a channel on a head to a well in the labware. **E**. Scheduling sets a flow variable for each piston/position pair, connecting aspirate and dispense wells to real-world transfers. The value of the flow variable represents the transfer volume. The wells and channels are color coded to demonstrate how a 60 *μ*l flow moves liquid between source and target wells. A complete schedule sets volumes for all flow variables, including many flows set to zero.

Each channel is constrained by a maximum liquid capacity, and multichannel instruments often have channels with identical capacities. The maximum liquid capacity of an instrument that uses disposable pipette tips is set by the tip volume. Pistons are defined by three parameters that control how liquid can move in and out of the attached channels:

1. The *minimum shot volume* that the piston can aspirate or dispense. Some pistons cannot expel liquid droplets below this threshold, and other instruments restrict the smallest dispense size due to increased error at small volumes.
2. The *maximum shot volume* that the piston can aspirate or dispense in a single operation. This volume may be different from the channel volume. For example, a pipettor’s piston may have a maximum shot volume of 1000 *μ*l even if it is attached to a channel with a 200 *μ*l tip; in this case, Pourfecto prevents the piston from aspirating more than 200 *μ*l.
3. A *shot size* for liquid handlers that dispense discrete droplets of a fixed size or handlers that limit the precision of their transfer volumes. For example, a continuous flow instrument can be programmed to dispense 1.24 *μ*l, while a discrete droplet instrument with a 0.1 *μ*l shot size must round its dispense to either 1.2 or 1.3 *μ*l.

The head definition connects channels and pistons with a binary mask **H**. Each element *h*_*pc*_ in **H** is 1 when piston *p* is connected to channel *c* and is 0 otherwise. Forcing a piston to aspirate causes all connected channels to aspirate the same volume. For example, setting the single piston on an eight channel pipette to aspirate 50 *μ*l places 50 *μ*l into all eight channels.

An instrument’s **deck** is a platform with slots that hold labware. Each slot can be accessed by the head and can hold compatible labware. Some robots have reconfigurable decks with movable slots for plates, tubes, bottles, and other equipment. Scheduling assumes that the instruments have been locked into a single configuration, i.e., the slot positions and labware types do not change during liquid handling. We refer to an instrument’s *configuration* as a pairing between a head and a deck layout. It is possible to schedule operations on a reconfigurable robot by defining separate liquid handlers for each configuration, although users need to account for the time to change between configurations. A single channel pipette has the most flexible deck with an infinite number of slots that can hold any labware. A human scientist’s benchtop can be viewed as the deck of a manual pipette.

A head’s geometry determines what labware on the deck are accessible to each channel. These constraints are captured by a second mask **L** for every head/labware pair that defines a finite set of possible positions for the head over the labware (Figure 2b–d). For example, an eight-channel pipette in the vertical orientation can access a standard 96-well microplate in 12 different positions—one for each column. Along each column, the eight channels can fit in only one configuration along the eight rows, as moving the channels vertically would push at least one channel outside of the plate (Figure 2b). A four-channel head can access the same microplate in 60 possible positions. Like the eight-channel pipette, the instrument can be placed in any of the 12 columns, but the four channels can be placed in one of five vertical positions along a single column: rows A–D at one extreme and rows E–H at the other (Figure 2c). The single-channel pipette has a simple mask for any labware, with the set of accessible positions taking the same shape as the labware, and each position corresponding to a single well (Figure 2d).

Some instruments have different masks for aspirating and dispensing. For example, instruments like the ARI Cobra are contact aspirators that dip four needles into the source liquid. The needles are aligned along a column of a 96 well plate, so the top needle cannot aspirate from rows F–H since the other needles would hit the bottom skirt of the plate. However, Cobra dispenses with the needles above the plate (by firing droplets into the wells), so the top needle can dispense into rows F–H. Users must define the aspirate and dispense masks for every labware when defining a new instrument head for Pourfecto.

Scheduling finds a sequence of triplets (aspirate position, dispense position, and piston flows) that recreate the transfer volumes identified during planning. Figure 2e illustrates the schedule for transferring 60 *μ*l from eight source wells of a 96 well plate into eight target wells of a 384 well plate using an eight-channel pipette. The scheduler finds a position over the source plate and aspirates 60 *μ*l with the piston.

It connects this flow to a dispensing position in the target plate that ensures the source liquids enter the correct target wells. The number of possible flow variables is quite large, as there is a single flow variable for every combination of instrument, labware, aspirate position, dispense position, and piston. The goal of scheduling is to find flows that minimize the work of pipetting while matching the planned volume transfers. We must connect the flow variables *q*_*ad*_ between aspirate position *a* and dispense position *d* with the planned transfer volumes *v*_*st*_ between source stock *s* and target stock *t*. The connection takes the form of a mass balance

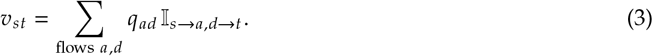

The indicator variable 𝕀 in the summand forces the total flow aspirated from the source stock *s* and dispensed into the target stock *t* to match the planned transfer volume from *s* to *t*. In practice, this indicator variable is replaced by the labware, head, and piston masks that define the connections between stocks during transfers. These connections are visualized in Figure 3 and can be interpreted as an operational view of scheduling. Volumes are routed through the labware masks into channels, then through the head mask into a piston, and finally through the position mask to an aspiration location. The scheduler connects the aspiration location to a dispense location, and the volumes are routed back through the masks into the target well.

**Figure 3:**
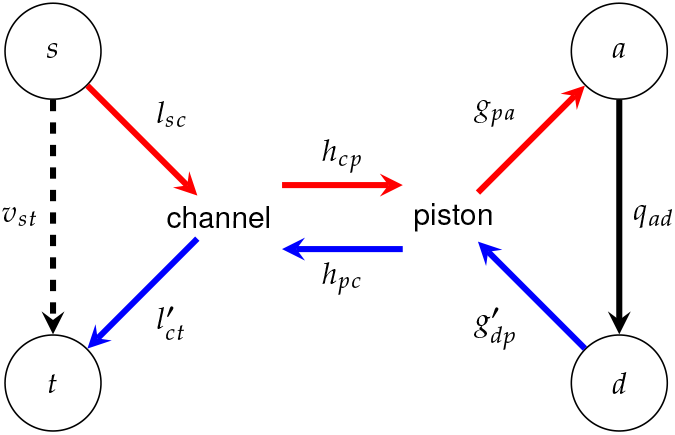
Masks for the labware (**L**), pipetting head (**H**), and piston geometry (**G**) connect a planned transfer volume *v* with a scheduled flow *q*, uniting the planning and scheduling views of liquid handling.

Mathematically, we can re-write Equation (3) using masks for the labware (**L**), head (**H**) and piston geometry (**G**) for both aspiration and dispensation:

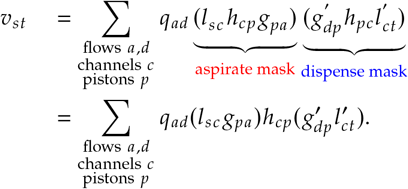

For computational efficiency, we combine the labware and piston masks when defining each robot in Pourfecto, and we note that the head mask is the same for both aspiration and dispensation (*h*_*cp*_ = *h*_*pc*_ and = 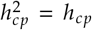. All of the masks can be precomputed once the robots are configured, so the above equations define a set of linear constraints that connect the planning and scheduling phases. We can use these equations to solve the planning and scheduling problems simultaneously across two stages. First, add the scheduling equations to the quadratic program used for planning (Equation (2)):

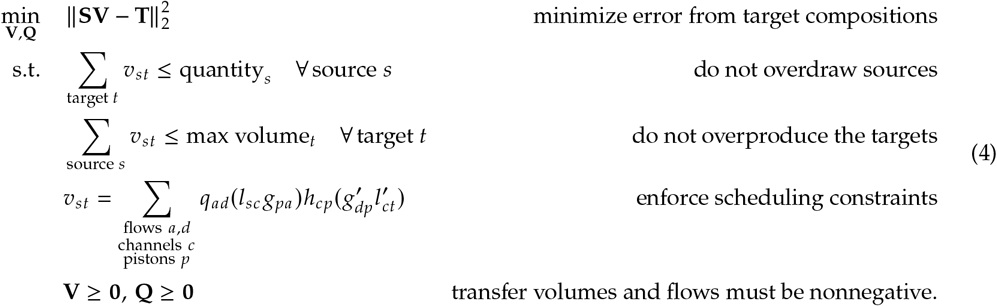

The above quadratic program finds a feasible schedule that minimizes the squared deviation from the target compositions. The feasible schedule may not optimally use the robots or minimize the operational cost of carrying out the liquid transfers. The second stage of scheduling solves an optimization problem that minimizes the operational cost without deviating from the original plan. The second optimization problem is

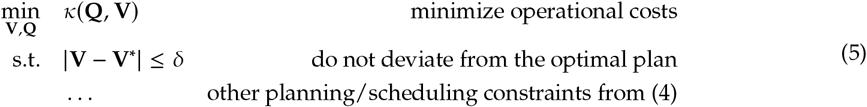

where **V**^*^ is the optimal planning solution calculated in the first scheduling stage (Equation (4)) and *δ* ≥ 0 is a threshold set by the user. Setting *δ* = 0 forces scheduling to adhere exactly to the optimal plan. Larger values of *δ* allow greater deviation from the target stock compositions to decrease the operational cost of liquid handling. The cost function *κ* encodes the user’s preferences for operational efficiency. This function could include the monetary costs of transfers (consumable tips or wear on hardware), the time required for a transfer, or the setup time for an instrument. By default, Pourfecto minimizes the total flow, which is a 1-norm approximation for minimizing the number of liquid transfers. This cost function is linear, so the scheduling problem becomes a quadratic problem (stage 1) followed by a linear program (stage 2) and can be efficiently solved for problems with hundreds or thousands of stocks. Other cost functions provide greater control over scheduling with the tradeoff of converting the stage 2 problem into a more computationally expensive mixed-integer linear program (MILP). Pourfecto’s other cost functions are described in Table 1.

**Table 1:** Pourfecto users can select multiple cost functions during phase 2 of scheduling (Equation (5)). The notation *expr* ⇒ 𝕀 defines a binary indicator variable I that takes the value 1 when *expr* is true and 0 otherwise.

| Objective | Cost function ($\kappa$) | Problem Type |
| --- | --- | --- |
| Minimize total flow with optional instrument-specific costs ($\omega_r$). This objective approximates the minimum number of flows objective (below) and is the default in Pourfecto. | $\kappa(\mathbf{Q}) = \sum_{\text{flows } a,d} \omega_r q_{ad}$ | LP |
| Minimize number of flows, which corresponds to the fewest movements and aspirations/dispenses. | $\kappa(\mathbf{Q}) = \sum_{\text{flows } a,d} \mathbb{I}_{q_{ad}}$<br>$q_{ad} > 0 \implies \mathbb{I}_{q_{ad}}$ | MILP |
| Minimize number of sources used. | $\kappa(\mathbf{V}) = \sum_s \mathbb{I}_s$<br>$\sum_{\text{sources } s} v_{st} > 0 \implies \mathbb{I}_s$ | MILP |
| Minimize number of labware used. | $\kappa(\mathbf{Q}) = \sum_l \mathbb{I}_l$<br>$\sum_{\text{flows } a,d} q_{ad} > 0 \implies \mathbb{I}_l$ | MILP |
| Minimize number of instruments (robots) used. | $\kappa(\mathbf{Q}) = \sum_r \mathbb{I}_r$<br>$\sum_{\text{labware } a,d \text{ on robot } r} q_{ad} > 0 \implies \mathbb{I}_r$ | MILP |

### 2.3 Slotting and compiling into machine instructions

Once an optimal schedule has been found, Pourfecto compiles the schedule into instructions for each instrument. For each instrument, the compiler identifies all source and target labware used by that instrument and searches for a configuration to slot the labware onto the instrument’s deck. Pourfecto slots using a greedy algorithm that places pairs of source and target labware until all slots are filled. Once a labware has been slotted, the compiler writes transfer instructions using each instrument’s file format. The slotting algorithm repeats until all the necessary source/target pairs have been placed on the instrument. Each iteration produces a run file for the instrument that is sent to the lab along with a set of slotting instructions for the technicians operating the instruments.

### 2.4 Implementation

Pourfecto is available as a Julia package at http://jensenlab.net/tools/. We implemented Pourfecto in Julia v1.8 using the JuMP mathematical modeling framework [17] and Gurobi optimizer v10.0.0 [18]. Pourfecto’s unit conversion features use the Unitful.jl package [19]. All benchmarking experiments were performed on a 4.05 GHz 8-core Apple M3 processor with 16 Gb of RAM. All code used to produce the experiments is available in the package repository.

Pourfecto can be configured to use new liquid handlers by specifying the following four components:

1. A head definition that includes channel/piston parameters and a head mask.
2. A deck definition indicating the number of slots and each slot’s labware compatibility.
3. A set of labware masks for each head.
4. A compiling function that converts a Pourfecto schedule into a run file for the instrument.

Our Julia implementation of Pourfecto includes methods that can be overloaded for new liquid handler definitions, including examples for a range of instruments. Writing an instrument’s compiling function is, in our experience, the most labor-intensive step in defining a liquid handler. Each liquid handler typically requires a unique (and sometimes proprietary) file format, with little standardization between instruments. If necessary, the protocols can be compiled into generic file formats (e.g., CSV files) and manually uploaded to the instrument’s control software.

## 3 Results

We tested Pourfecto using an example set of seven liquid handlers (Table 2). Each liquid handler has a unique head or deck design and a set of compatible labware.

**Table 2:** Pourfecto uses a common framework to plan and schedule across manual and robotic liquid handlers. Examples in this work use a single channel head for the NIMBUS, but other heads are available.

| Name | Manufacturer | Type | Channels | Pistons |
| --- | --- | --- | --- | --- |
| Single Channel | USA Scientific | Manual, Handheld | 1 | 1 |
| Eight Channel | Eppendorf | Manual, Handheld | 8 | 1 |
| NIMBUS™ | Hamilton | Robotic, Reconfigurable Deck | 1* | 1* |
| Cobra 4™ | ARI | Robotic, Fixed Deck | 4 | 4 |
| PlateMaster™ | Gilson | Manual, Fixed Deck | 96 | 1 |
| Mantis™ | Formulatrix | Robotic, Fixed Deck | 1 | 1 |
| Tempest™ | Formulatrix | Robotic, Fixed Deck | 8 | 8 |

### 3.1 Preparing combinatorial growth media

We used Pourfecto to generate optimal liquid handling protocols for preparing media for a high-throughput microbial growth experiment (Figure 4a). Our goal was to prepare 480 unique 200 *μ*l media containing unique concentrations of up to 48 reagents (plus water). The target wells were assigned to 96 well microplates (Figure 4a, labwares 4–8). Source stocks were located in three types of labware (labwares 1–3):

**Figure 4:**
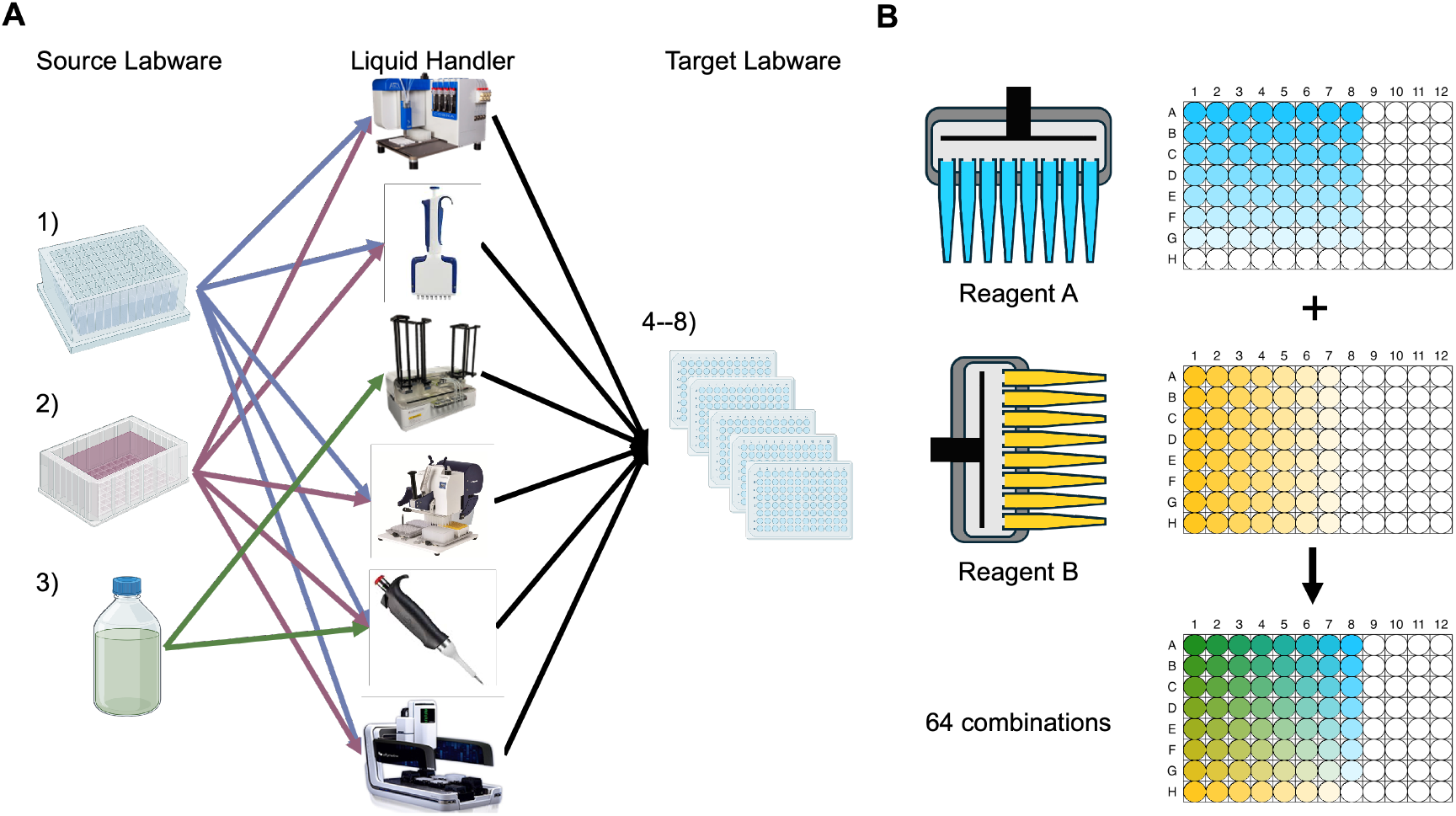
Pourfecto schedules liquid handling protocols using multiple liquid handlers. **A**. Pourfecto generated liquid handler protocols to create 480 microbial growth media containing unique combinations of up to 48 reagents. Each liquid handler has constraints that limit the compatible labware. The green bottle, for example, is only compatible with two of the six instruments. **B**. Pourfecto found the optimal 16-flow solution for preparing a reagent synergy experiment by selecting a multichannel pipette in two different orientations.

1. A deep 96-well plate containing the 48 reagent stocks and three stocks of water. (The remaining 45 wells were empty.)
2. An SLAS format reagent reservoir containing water.
3. A 1000 ml bottle containing water.

We allowed Pourfecto to use any combination of six liquid handlers to prepare the target stocks: an ARI Cobra, an eight-channel handheld pipette, a Formulatrix Tempest, a 96-channel Gilson PlateMaster, a single-channel pipette, and a Hamilton NIMBUS. As indicated in Figure 4a, not every labware was compatible with every liquid handler. The multiwell plate-based instruments, for example, cannot hold a 1000 ml bottle on their deck, and plate fillers like the Tempest cannot accommodate multiwell plates as sources. Pourfecto satisfies these constraints as it searches for solutions.

Pourfecto found a schedule that matched every reagent target for all 480 media within a tolerance of 0.1%. The schedule used three of the five instruments (Table 3). The ARI Cobra was used to dispense from the deep well plate to five 96 well target plates. The Cobra holds one source and one target plate at a time, so Pourfecto slotted the target plates in five batches and generated separate run files for each batch. The three wells of water in the deep well plate were not sufficient to fill all of the target wells. Pourfecto used both the 96-channel Gilson PlateMaster and a multichannel pipette with the water reservoir to complete the water dispensing. It did not use any water from the 1000 ml bottle.

**Table 3:** Pourfecto modifies schedules in response to changing inputs. Removing the water reservoir as a source forced Pourfecto to use the water bottle and switch from plate-compatible instruments to the Formulatrix Tempest. The second schedule follows an identical plan but requires a schedule with a higher operational cost.

| Instrument | Transfer Volume (ml) |  |
| --- | --- | --- |
|  | With Reservoir | Without Reservoir |
| ARI Cobra™ | 1.767 | 8.727 |
| eight channel pipette | 0.216 | 0 |
| Formulatrix Tempest™ | 0 | 87.271 |
| Gilson PlateMaster™ | 94.003 | 0 |
| single channel pipette | 0 | 0 |
| Hamilton NIMBUS™ | 0 | 0 |
| **Total Volume** | 95.990 | 95.990 |
| **Total Flow** | 2.779 | 95.990 |

Pourfecto’s schedule found the minimum number of well-to-well transfers—18,089—to complete the protocol. Pourfecto selected the PlateMaster and the multichannel pipette to complete many transfers simultaneously with a single piston movement. Using other instruments, such as a single channel pipette or the NIMBUS, would have required more flow operations to complete the same set of transfers.

In a follow-up experiment, we removed the reagent reservoir and rescheduled the same target wells, thereby forcing Pourfecto to use the 1000 ml bottle as a water source. The PlateMaster and the multichannel pipette cannot aspirate from the 1000 ml bottle, so Pourfecto instead used the Tempest to dispense water. The new schedule had the same planning accuracy but required 34 times more total flow (Table 3). Pourfecto’s flexibility when scheduling around multiple real-world constraints allows it to find solutions to complex liquid handling tasks.

### 3.2 A minimial pipetting protocol for a reagent synergy experiment

As a second test, we instructed Pourfecto to prepare a protocol for a pairwise reagent synergy experiment. Such assays commonly use a “checkerboard” layout that varies the concentration of each reagent along a microplate’s rows and columns [20], [21](Figure 4b). The concentration of reagent A (blue) decreases linearly along each row, and reagent B (yellow) decreases linearly across each column. Together, the reagents form 64 unique combinations. A scientist can quickly plan and schedule this experiment with an eight-channel pipette. Reagent A can be dispensed in each row with the pipette in a horizontal orientation, and reagent B can be dispensed with the pipette in a vertical orientation. Using the multichannel pipette is the optimal solution and creates the 64 combinations using 16 operations. The 96 channel pipette cannot be used because it cannot dispense volumes independently by row or column, and a single channel pipette would require 2 × 64 = 128 operations.

We verified that Pourfecto finds the optimal solution for the checkerboard assays. To ensure that it found the minimum pipetting solution, we instructed it to use the secondary objective that minimizes the total number of active flow operations. Pourfecto found and scheduled the 16-operation multichannel protocol in 1.45 seconds. This experiment demonstrates how Pourfecto can be used to optimize manual pipetting plans and find solutions that match human expectations.

### 3.3 Performance Benchmarking

We benchmarked Pourfecto’s scalability by creating variations of the growth media experiment from section 3.1. For each experiment, we varied the number of media targets from 96–480, the total number of wells from 192–577 (including the source labware), and the number of unique reagents in the media from 12–48. We used Pourfecto to find 496 liquid handling plans using one to five instruments. Every experiment included the single-channel pipette (to ensure that every plan was feasible) and a random selection of the remaining instruments. All protocols matched every reagent target for every stock within the numerical tolerance of the solver. The worst case runtime was 5 minutes 30 seconds for a plan that included 577 wells (97 sources and 480 targets), 36 reagents, and all 5 instruments (a total of 8 pistons). We profiled Pourfecto and found that the runtime of the algorithm is dominated by writing constraints to set up the optimization problem (Supplementary Material 2). In particular, writing the scheduling constraints was the rate-limiting step. This constraint is written in two stages. The first stage pre-computes the aspirate and dispense nodes for every well, which in the worst case scales with 𝒪 (*wp*) time complexity, where *w* and *p* are the number of wells and pistons in the problem, respectively. The second stage writes the constraints by iterating over each pair of wells and then iterating over each flow/well pair. The time complexity of this second stage is approximately 𝒪(*w*^2^ (*wp*) ^2^) = 𝒪 (*w*^4^*p*^2^). Using a linear model, we empirically verified that the total runtime of Pourfecto scales with the predicted time complexity. Interestingly, the number of reagents considered in the plan has little effect on the runtime, so Pourfecto is well-suited to planning experiments with a large number of reagents.

## 4 Discussion

No single piece of software covers all of the operations in an autonomous lab. State-of-the art tools provide a subset of the following four features:

1. **Protocol Planning and Scheduling:** Generating lab operations from abstract inputs such as experimental designs or reagent requests.
2. **Protocol Compiling:** transforming a set of lab operations into executable code for instruments.
3. **Lab Orchestration:** Coordinating when and where laboratory processes are executed. Lab orchestration problems include traditional scheduling frameworks such as job shops and flow shops.
4. **Multi-instrument Coordination:** Splitting a single protocol across multiple instruments. Schedulers that support multi-instrument scheduling offer more flexibility compared to single-instrument schedulers.

Most commercial software specializes in lab orchestration to drive integrated platforms with multiple (simultaneous) users and human-designed processes[12], [13], [22]–[24] (Table 4). Other tools allow scientists or automation engineers to create protocols using no-code interfaces to map out workflows[11], [14], [25]. Once a protocol has been defined, these software packages compile the protocol and place it into a managed queue of lab tasks. Other tools focus on scheduling problems such as deciding the sequence of protocols to be executed on a given instrument or minimizing the movement of robotic arms while handling labware[26], [27].

**Table 4:**
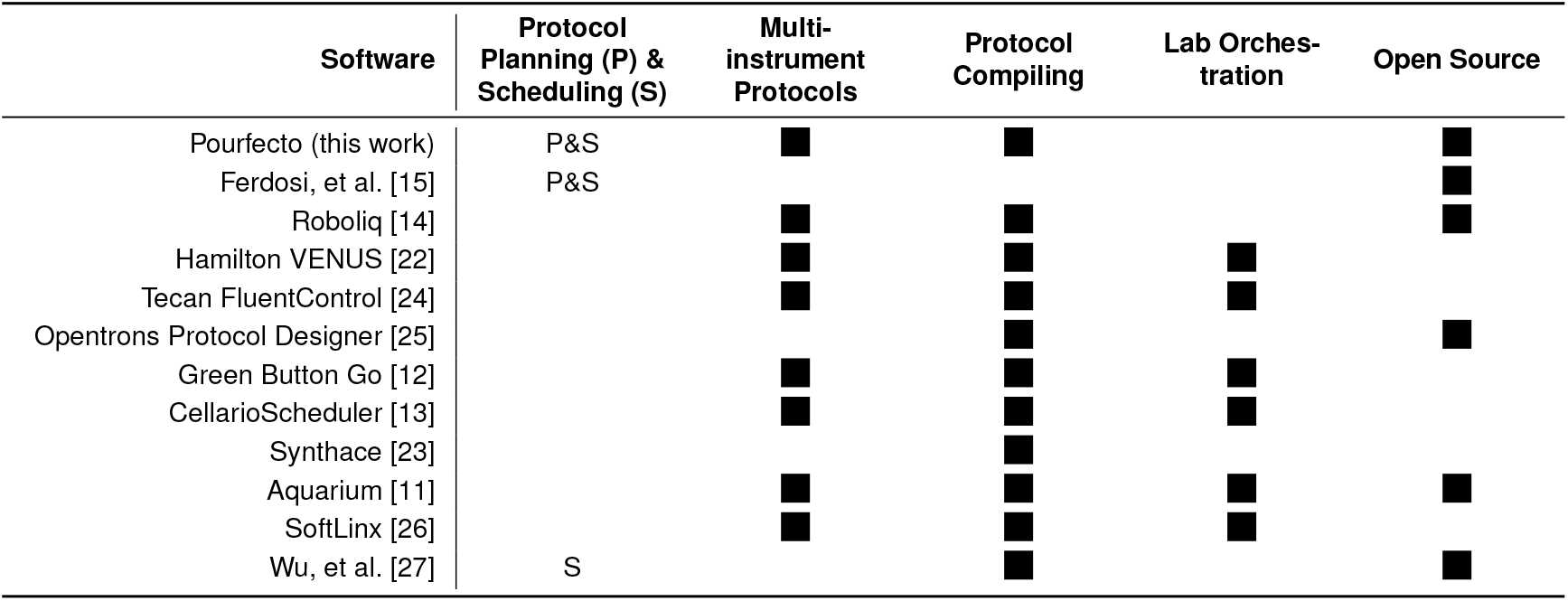
Pourfecto plans and schedules protocols across multiple instruments, whereas many existing solutions require human defined protocols and focus on integrating multiple instruments with a common user interface.

| Software | Protocol Planning (P) & Scheduling (S) | Multi-instrument Protocols | Protocol Compiling | Lab Orchestration | Open Source |
| --- | --- | --- | --- | --- | --- |
| Pourfecto (this work) | P&S | ■ | ■ |  | ■ |
| Ferdosi, et al. [15] | P&S |  |  |  | ■ |
| Roboliq [14] |  | ■ | ■ |  | ■ |
| Hamilton VENUS [22] |  | ■ | ■ | ■ |  |
| Tecan FluentControl [24] |  | ■ | ■ | ■ |  |
| Opentrons Protocol Designer [25] |  |  | ■ |  | ■ |
| Green Button Go [12] |  | ■ | ■ | ■ |  |
| CellarioScheduler [13] |  | ■ | ■ | ■ |  |
| Synthace [23] |  |  | ■ |  |  |
| Aquarium [11] |  | ■ | ■ | ■ | ■ |
| SoftLinX [26] |  | ■ | ■ | ■ |  |
| Wu, et al. [27] | S |  | ■ |  | ■ |

Pourfecto brings planning and scheduling capabilities to multi-instrument laboratories. By automating protocol creation, Pourfecto acts as a layer of abstraction between scientists who provide experimental designs and the engineers who configure automated laboratories. Compared to existing solutions for protocol scheduling, Pourfecto can leverage the capabilities of different instruments and satisfy the logistical constraints that arise when splitting protocols across robots and human technicians. For example, Pourfecto found protocols for the microbial growth media experiment that correctly used multichannel instruments for dispensing bulk-filled reagents and used other instruments for combinatorial mixing steps. These instructions were not given to the software; the plans were instead found by the solver by minimizing the number of operations in the protocols. Pourfecto computes protocols on demand, allowing scheduling to happen just prior to the experiment’s execution. If, for example, an instrument goes offline or a reagent stock runs out, Pourfecto can compute a new plan that uses other resources in the lab, allowing laboratories to adapt to unforeseen events and create new protocols on the fly.

Just-in-time protocol design requires Perfecto to find solutions in seconds or minutes. We chose a linear/mixed-integer programming framework over other optimization frameworks, such as stochastic search algorithms, because it finds provably optimal solutions, scales well to large problems (with multiple types of liquid handlers, reagents, and labware) and is compatible with the objectives and constraints we identified. Pourfecto outputs clear metrics of solution quality, such as how closely its protocols produce target stocks. It can identify infeasible problems and provide interpretable errors. For example, if Pourfecto detects an insufficient quantity of a source reagent, it instructs the user to increase the available volume of the reagent.

Our formulation of the planning and scheduling problems assumes that all pipetting is done in a single pass. Target wells cannot be used as sources for other targets. Pourfecto cannot, for example, plan and schedule serial dilution experiments on its own. Instead, a larger combinatorial optimization problem must be solved to identify the optimal intermediate stocks. For example, a rollout algorithm or another reinforcement learning technique can be used to optimize a sequential pipetting problem. Once a set of intermediate stocks is defined, Pourfecto can be used to solve the planning and scheduling subproblem and return a reward signal to the outer problem. Here again, Pourfecto’s speed and determinism make it an ideal part of a lab orchestration pipeline.

Pourfecto requires users to specify the operational objectives that best suit the needs of their automation platform. Users need to decide, for example, whether they prioritize solutions that minimize pipetting time/costs over the need to prepare larger quantities of reagents. Aligning objectives to real-world measures of cost is not always clear, and user preferences can change from experiment to experiment. One option is to run Pourfecto in parallel with multiple objectives and evaluate candidate protocols, creating a “Pareto front” of optimal solutions. As mentioned in Section 2.4, users can also define custom objectives that align with their cost measures.

Pourfecto does not consider consumable usage, e.g. disposable tip costs, for contact-based liquid handlers. Engineers often schedule protocols that minimize tip usage by finding protocols that change tips less frequently. Our definition of liquid handlers does not include a measure of tip usage, so tip-based objectives cannot be tracked by Pourfecto. Some downstream instrument-level schedulers, however, do find minimum-tip paths when planning the physical liquid handler movements for a given protocol[22]. While the choice of protocol bounds tip usage, the protocols designed by Pourfecto can be further optimized to reduce consumables. Future versions of Pourfecto could include tip usage as part of the cost framework. Designing such a feature as an optional extension would reduce the need to add constraints to the problem if the user is not interested in those objectives.

Automation plays an increasingly important role in science. Cloud labs, self-driving labs, and high-throughput screening facilities are expanding to include new types of instruments, and we believe Pourfecto’s emphasis on multi-instrument scheduling fills an important gap in the automation software landscape. Pourfecto can be combined with other schedulers that optimize plate layouts, maintain reagent stocks, and coordinate lab tasks to facilitate the shift toward scientific autonomy. Pourfecto uses linear, quadratic, and mixed-integer programming as heuristic approaches for finding optimal protocols. Other approaches, such as dynamic programming, reinforcement learning, simulated annealing, and genetic algorithms, are also valuable tools for tackling automation problems. Looking ahead, large language models are promising tools for parsing experimental designs and scheduling protocols [28], and future automation software may combine artificial intelligence with traditional deterministic optimization. Work in all of these areas will remove barriers between planning abstract experimental designs and scheduling physical laboratory processes.

## 5 Acknowledgements

This work was supported by NSF Award #2339026. BMD was supported in part by the SLAS Graduate Education Fellowship and the University of Michigan Rackham Predoctoral Fellowship.

## Footnotes

† Our piston/channel framework for liquid handlers uses terms from the design of standard micropipettes. Many liquid handling robots use mechanisms other than piston displacement, such as diaphragms, solenoids, piezoelectrics, ultrasound, or electrowetting to aspirate and dispense; the exact pipetting mechanism does not matter, and Pourfecto can schedule all of these devices. Similarly, the virtual channels in our framework could be a physical channel in a tip based-dispenser or a nozzle, needle, or tube on other instruments.

