## Supplementary Materials for "Planning and scheduling biological experiments across multiple liquid handling robots"


#### 1 Reagent Precedence

By default, Pourfecto is prohibited from using sources with reagents that are not in the target stock. For example, when planning to make the target

$$\mathbf{t} = \begin{pmatrix} 10 & \text{mg glucose} \\ 5 & \text{mg ampicillin} \\ 0 & \mu\text{l ethanol} \\ 1000 & \mu\text{l water} \end{pmatrix}$$

with the sources

$$\mathbf{s}_1 = \begin{pmatrix} 16 & \text{g/l glucose} \\ 0 & \text{g/l ampicillin} \\ 0 & \text{\% volume ethanol} \\ 100 & \text{\% volume water} \end{pmatrix}, \quad \mathbf{s}_2 = \begin{pmatrix} 0 & \text{g/l glucose} \\ 2 & \text{g/l ampicillin} \\ 0 & \text{\% volume ethanol} \\ 100 & \text{\% volume water} \end{pmatrix}, \quad \text{and} \quad \mathbf{s}_3 = \begin{pmatrix} 0 & \text{g/l glucose} \\ 20 & \text{g/l ampicillin} \\ 100 & \text{\% volume ethanol} \\ 0 & \text{\% volume water} \end{pmatrix},$$

Pourfecto is blocked from using source $\mathbf{s}_3$ because it contains ethanol, while the target $\mathbf{t}$ does not. However, it is often impossible to match every target reagent concentration with the available sources. In the above example, the concentration of ampicillin in stock $\mathbf{s}_2$ is not high enough to match the target concentration of 5 mg, so Pourfecto's default solution under-delivers glucose and ampicillin to keep the final stock volume at 1000 $\mu$l (Table 1). This is one of many sub-optimal solutions to this unsatisfiable problem, and users might value other tradeoffs in selecting a solution. One option is to prioritize matching the glucose target over ampicillin (or vice versa). Another option is to let Pourfecto use source $\mathbf{s}_3$ because it contains a higher concentration of ampicillin, even though the final stock would contain ethanol. Pourfecto cannot make these decisions on its own, so we included a mechanism for users to specify their preferences.

Pourfecto's optional input, called *precedence*, establishes priority between reagents. In the precedence system, each reagent is assigned a non-negative integer. For a pair of reagents, the reagent with the lower precedence is given higher priority. The precedence system allows for duplicate precedence levels. By default, all reagents in the target stock are assigned to precedence level 1, indicating that they should all be handled equally by Pourfecto. Superfluous reagents are assigned to precedence level 0 to indicate that avoiding them is more important than matching target reagents. This default precedence scheme leads to the first solution in Table 1, where Pourfecto balances under-delivering glucose and ampicillin to avoid including ethanol.

Solvents are a common use case for precedence. For example, many small molecules are poorly soluble in water but highly soluble in DMSO. Users must choose between adding a small amount of a concentrated stock with DMSO or a large volume of dilute stock with water. The tradeoff depends on how sensitive an experiment is to unwanted reagents and how closely the target quantities must be matched.

Setting water to precedence level $\infty$ (or more accurately $2^{64} - 1$ as a 64-bit unsigned integer) instructs Pourfecto that water is the least important reagent in the plan. The second and third rows of Table 1 highlight the impact of other precedence schemes on Pourfecto's solution. Setting ampicillin to precedence level 2 instructs Pourfecto to first hit the glucose target and then try to hit the ampicillin target. The third solution keeps glucose and ampicillin at precedence level 1, but now allows Pourfecto to use ethanol in its solution

| Precedence | | $\mathbf{v}^*$ | $\mathbf{t}^*$ | |
|---|---|---|---|---|
| glucose<br>ampicillin<br>ethanol<br>water | 1<br>1<br>0<br>$\infty$ | $\begin{pmatrix} 592.25\ \mu\text{l} & \mathbf{s}_1 \\ 407.75\ \mu\text{l} & \mathbf{s}_2 \\ 0\ \mu\text{l} & \mathbf{s}_3 \end{pmatrix}$ | $\begin{pmatrix} 9.48 & \text{mg glucose} \\ 0.81 & \text{mg ampicillin} \\ 0 & \mu\text{l ethanol} \\ 1000 & \mu\text{l water} \end{pmatrix}$ | Under the default settings, $\mathbf{s}_3$ is not used, and neither target is matched. |
| glucose<br>ampicillin<br>ethanol<br>water | 1<br>2<br>0<br>$\infty$ | $\begin{pmatrix} 624.94\ \mu\text{l} & \mathbf{s}_1 \\ 375.06\ \mu\text{l} & \mathbf{s}_2 \\ 0\ \mu\text{l} & \mathbf{s}_3 \end{pmatrix}$ | $\begin{pmatrix} 10 & \text{mg glucose} \\ 0.75 & \text{mg ampicillin} \\ 0 & \mu\text{l ethanol} \\ 1000 & \mu\text{l water} \end{pmatrix}$ | Glucose is prioritized over ampicillin. |
| glucose<br>ampicillin<br>ethanol<br>water | 1<br>1<br>2<br>$\infty$ | $\begin{pmatrix} 624.94\ \mu\text{l} & \mathbf{s}_1 \\ 138.99\ \mu\text{l} & \mathbf{s}_2 \\ 236.08\ \mu\text{l} & \mathbf{s}_3 \end{pmatrix}$ | $\begin{pmatrix} 10 & \text{mg glucose} \\ 5 & \text{mg ampicillin} \\ 236.08 & \mu\text{l ethanol} \\ 763.93 & \mu\text{l water} \end{pmatrix}$ | Ethanol is minimized only after matching the glucose and ampicillin targets. |

**TABLE 1:** Pourfecto's precedence system helps users balance solution tradeoffs. By default, Pourfecto prohibits solutions that use stocks with unwanted reagents. However, users can relax this constraint for infeasible problems to find sub-optimal solutions. Users can also specify if certain reagents should be avoiding with a higher priority.

by setting it to precedence level 2. Ethanol is placed at a lower precedence level than water to ensure that it is still minimized.

Precedence requires a small change to the formulation of the planning problem (Equation 1). To facilitate precedence constraints, the planning problem is solved sequentially by iterating through each precedence level. If there are $P$ unique precedence levels, the planning problem is solved $P$ times. For a particular precedence level $p$, we create a vector of indicator variables $\boldsymbol{\phi}^p$. The indicator $\phi_r^p$ is equal to one if reagent $r$'s precedence level is less than or equal to $p$, and zero otherwise. The vector $\boldsymbol{\phi}^p$ appears in the objective of the planning problem, focusing the objective only on the reagents at or below the current precedence $p$. After Pourfecto finds a solution for precedence level $p$, a constraint is added to fix the quantities of all reagents at or below precedence level $p$ before finding a solution for precedence level $p+1$ via the following optimization problem:

$$\begin{aligned}
\min_{\mathbf{V}} \quad & \left\| \boldsymbol{\phi}^p \cdot (\mathbf{SV} - \mathbf{T}) \right\|_2^2 && \text{minimize error from target compositions} \\
\text{s.t.} \quad & \mathbf{V} \geq \mathbf{0} && \text{transfer volumes must be nonnegative} \\
& \sum_{\text{target } t} v_{st} \leq \text{quantity}_s \quad \forall \text{source } s && \text{do not overdraw sources} \\
& \sum_{\text{source } s} v_{st} \leq \text{max volume}_t \quad \forall \text{target } t && \text{do not overproduce the targets.} \\
& \boldsymbol{\phi}^k \cdot \mathbf{S} |\mathbf{V} - \mathbf{V}^*| \leq \delta \qquad \forall k < p && \text{do not deviate from earlier precedence levels} \\
& \text{precedence}_r \leq p \implies \phi_r^p \quad \forall \text{reagent } r && \text{add level } p \text{ reagents to } \boldsymbol{\phi}^p.
\end{aligned} \tag{1}$$

### 2 Pseudocode for Pourfecto (without precedence)

**Data:** Source Wells ($sources$) , Target Wells($targets$), Instrument Configurations ($instruments$), operation cost function ($\kappa$)
**Result:** A set of well-to-well transfers for every source labware, target labware, and instrument paring ($transfers$)

```
labware= the set of all unique labware that contain every unique source and target well;
L = length(labware);
I = length(instruments);
W = length(union(sources,targets));
Construct matrices S and T, containing the source concentrations of each reagent and the final target quantities for all wells, respectively ;
Initialize empty vectors for aspirate nodes (asp) and dispense nodes(disp);
for l ∈ 1 : L do
    for i ∈ 1 : I do
        compute all aspirate and dispense nodes for instruments_i and labware_l and add them to asp and disp;
begin model
    define V ∈ R_+^{W×W}, the set of well to well transfers;
    define Q ∈ R_+^{A×D}, the set of all flows between aspirate and dispense nodes;
    add target well capacity and source overdraft constraints;
    add constraint to check that entries in Q use the same instrument and piston;
    for w_s ∈ 1 : W do
        for w_t ∈ 1 : W do
            initialize connections;
            for a ∈ asp_{w_s} do
                for d ∈ disp_{w_t} do
                    if w_s, w_t connected through node pair a,d then
                        add (a, d) to connections;
            add constraint V_{w_s,w_t} = Σ_connections q_{a,d};
    minimize ||SV − T||_2^2;
    add constraints |V − V*| ≤ δ ;
    minimize κ(Q, V);
return optimal plan V* and schedule Q*
```
